# Mark-Release-Recapture Experiments in a North Sumatra Oil Palm Plantation: Assessing the Impact of Super-Male Planting Material on Fruit Set and *Elaeidobius kamerunicus* population size

**DOI:** 10.64898/2026.09.10.750641

**Authors:** Yves Dumont, Laurence Beaudoin-Ollivier, Hervé Rey, Axel Labeyrie, Camille Madec, Anna Doizy, Indra Syahputra, Dadang Afandi, Florence Jacob

**Author notes:** Correspondence: Yves Dumont.

## Abstract

Yield improvement is a main issue in the oil palm industry. Several ways to achieve this goal are possible, like, for instance, improving fruit set. Here we explore a planting strategy to increase two major factors influencing pollination; i.e. (1) the number of individuals of its main pollinator, *Elaeidobius kamerunicus Faust*, and (2) pollen availability, by mixing oil palm planting material, usually highly feminine, with a new oil palm material, the “super-male” (SM) planting material, with a high ratio of male inflorescences. We design and study a field trial comprising extremely feminine planting material mixed with different SM densities. Using Mark-Release-Recapture experiments we estimate the average dispersal of the pollinators over three different periods of the year, from 2017 till 2022. Second, using phenological data recorded over the same period, we studied the impact of the SM planting density on the population size of the pollinators. Finally, we assess the impact over time of different SM planting densities on the fruit set, the fruit-to-bunch, and the number of trapped wild pollinators. The combination of SM material with a highly feminine planting material provides very promising results in terms of fruit set, linked with increased pollinator population and pollen availability, demonstrating its potential for application in oil palm plantations.

## Introduction

Oil palm industry is one of the main industries in Indonesia, in particular in North-Sumatra. However, it is only in 1911 that oil palm cultivation started in Sumatra, initiated by the Franco-Belgium company Socfin (“Société financière des Caoutchoucs”). Since then, oil palm cultivation has developed with the objective of improving oil yield per hectare. This production increase was made possible by breeding, improving cultivation practices, and finally improving pollination^1^. Indeed, the optimum yield can be reached with a fruit set ranging from 60 to 80%, with two critical factors involved: pollen availability and pollen transport.

Two types of pollination occur in oil palm, which is a monoecious species with alternating cycles of male and female inflorescences on the same plant: entomophilic pollination^2^ and anemophilic pollination^3^. The second one is particularly effective when oil palms are tall. The entomophilic pollination mainly occurs when planted oil palms are young (below 8-10 years old).

Introduced from Africa in Malaysia in 1980, and in Indonesia in 1982, the pollinating weevil *E. kamerunicus* Faust 1898 (Curculionidae: Curculioninae: Derelomini) has ensured since then most of the entomophilic pollination in South-East Asian oil palm plantations. Before its introduction, pollination was done manually and required a large taskforce, which was very costly. A review about *E. kamerunicus* can be found in ^4^.

Other oil palm pollinators exist^5^, like *E. subvittatus* and *E. plagiatus*^2^, the last one being, like *E. kamerunicus*, a good pollinator^6^. However, *E. kamerunicus* was chosen due to its high-pollen carrying capacity^7–9^, and thus is considered the most effective for pollination. That is why it has been the only one to be introduced in Indonesia and Malaysia in the early-1980s^10,11^.

*E. kamerunicus* is well known as being one of the main pollinators of oil palms, specifically in tropical regions like West and Central Africa, from where it is native^2^. Oil palm pollination is an example of mutualistic interaction between a crop and its insect pollinators: *E. kamerunicus* complete its whole life cycle on male inflorescences, with several successive immature stages that are easy to identify: egg, larva, prepupa, pupa and then imago. From eggs deposit till the emergence of imagos, it takes, on average, between 8 and 12 days^12^. Attracted by volatile compounds (aniseed scent) emitted by male inflorescences in anthesis, the female deposit one egg per flower and, at the same time, loads with pollen. On average a female can lay 200 eggs^12^ in anthesizing male inflorescences. Then, when the neonate larva hatches, the flower serves as food for the larva. In the lowland of North-Sumatra, males live longer (52 days) than females (38 days)^12^. After visiting a male inflorescence in flower, weevils, loaded with pollen, continue to seek out new male inflorescences in anthesis in order to feed, reproduce and, in the case of the females, lay their eggs. However, female inflorescences in anthesis also emit volatile compounds, similar to those of the male inflorescences, such that the deceived *E. kamerunicus* may visit the female inflorescences, feed on nectar, and thus release the pollen, contributing to the pollination of female flowers and thus to the fruit set. In theory, the higher the number of visits of (viable) pollen carrying weevils (during the anthesis stage that only lasts two (five) days for female (male) inflorescences^1^), the better the fruit set, and, thus, the fruit yield.

Since the introduction of *Elaeidobius kamerunicus* in South-East Asia with great success^10^, several works tried to estimate the density of pollinators, per hectare, required to reach a certain fruit set (see for instance ^13–15^ and references therein). However, so far, the most precise results were obtained in^16^, where the authors estimated the weevil’s population per hectare and the related fruit set, and, also, the fruit set versus the male inflorescences per hectare and per month. In particular, if the number of male inflorescences in anthesis per hectare and per month is below 50, then the fruit set is poor (see also ^1^, page 132, Figure 5.17).

Several factors may influence the weevil’s population along the year^7,8,17^: rainfall, natural predators such as rats, palm cultivar (size and number of male inflorescences), and, of course, the density of anthesizing male inflorescences per hectare, leading to considerable variations in fruit set of standard plantations. At the same time, pollen availability can be influenced by many factors itself. Here we focus on the oil palm sex ratio, which is determined by environmental and genetic factors^1^, as a critical parameter linked with the number of male inflorescences produced. For example, a ratio of 90% of female inflorescences was reported in one oil palm planting in Malaysia, and such ratio leads to a low density of male inflorescences associated with a fruit set deficit, and thus to a yield loss^1^. It is common to observe very high female to male inflorescence ratios in plantations, over short or longer periods, and this can be linked with diverse factors such as very favorable environmental conditions, age of the palms, seasonal variations, and type of cultivar. Indeed, breeding efforts to increase yield have resulted in commercial cultivars with increased bunch index, which implies a higher number of female inflorescences at the expense of the male inflorescences. This can exacerbate pollination issues in commercial plantations, especially during the first years of harvest^8^.

To counterbalance these pollinations issues, planters have tried several approaches. While a standard commercial plantation density ranges between 128 and 143 palms per hectare, high-density planting have been considered in order to increase the competition between palms, inducing stress, and, thus, increasing the production of male inflorescences^18,19^, but, at the same time, decreasing the number of female inflorescences, and thus the yield. Other cultivars, based on genetic selection, have also been considered but without substantial improvements^17^. A novel approach, developed by PalmElit SAS, and based on the synergistic relationship between male inflorescences and *E. kamerunicus*, is to increase the male inflorescence density per hectare by adding super-male (SM) palms that are able to continuously produce a high number of male inflorescences. This approach is called the SM approach and will be developed later in the manuscript. Additionally, super-male palms may also be used for pollen production and harvest in support of assisted pollination.

*E. kamerunicus* pollen dispersal ability is another key parameter. It can be useful to develop new mathematical models to optimize the pollination strategies in commercial plantations, such as planting designs with “pollinator palms”. Dispersal ability over a certain period has been studied in terms of establishing population or in Hatch and Carry experiments^20^ showing a fruit set improvement within 200 m from the location of the release boxes.

However, so far, there are no pollen and/or pollinator dispersal results inside oil palm plots, while this knowledge is of main importance. Indeed, when considering the use of different types of plant material, particularly where one of these types is expected to produce significantly more male inflorescences – as is the case with SM palms – it is necessary to ensure that the viable pollen produced by the male inflorescences of the SM palms is dispersed by pollinators to the female inflorescences of the predominantly female cultivar. Then, knowing, at least, the average dispersal distance may be useful to optimize a planting design involving SM palms spread within productive oil palms.

Estimates of pollen dispersal ability and population size can be obtained using Mark-Release-Recapture (MRR) experiments. MRR experiments have been used by ecologists and entomologists for more than a century for a wide range of animals and insects, and the method was first described in 1896^21^. MRR consists of either capturing or bringing individuals (from another field), marking these individuals such that they can be identified, and then releasing them (back) into the targeted population. Later, recaptures occur, using traps, and marked individuals are counted within the samples^21^. MRR can be used to estimate various parameters, like the lifespan, the dispersal, etc, and, also, to obtain estimates of absolute population size^22^. It may provide reliable results as long as some basic assumptions are met. In particular, it must be assumed that marking has no effect on behavior and thus on capture probability compared to non-marked insects. Unfortunately, these MRR experiments, while useful, can be laborious, tedious to conduct, and also costly. Several improvements have been made, or are under study, to simplify the protocols and the analysis^23^.

In this work, we aim to provide new results related to *E. kamerunicus* average dispersal distance, population estimates, and to the impact of the SM treatment, i.e. SM plantation density, on the pollination efficiency, including the average fruit set and fruit to bunch.

## Materials & Methods

### The Mark-Release-Recapture-Experiments

We set up several series of the Mark-Release-Recapture experiments from February 2017 till December 2022. The MRR experiments were done at three different periods, representing dry or rainy periods, each year: February-March (Feb), June-July (Jun) and November-December (Nov). In North-Sumatra, the dry season lasts from May till September, the rainy season from October till April, with heavy rainfalls in November and December.

For each batch of release, we selected thousands (around 5000 individuals) weevils from Hatch & Carry cages containing male inflorescences collected in old plantations, and marked them using luminous powders: red (BioQuip product, Inc, USA: 1162R), yellow (BioQuip product, Inc, USA: 1162Y), orange (Sennelier, France: 648), and emerald green substitute (Sennelier, France: 869), one for each day of release. Marked insects were released in the center of an 8-hectare block (block 1) of the Super-Male trial (described in the section below). The releases took place over a maximum of five consecutive days, between 8.30 a.m. and 10.00 a.m., during maximum flight activity of the insects.

Before the start of the MRR experiments, a survey was conducted in the field to record and count the inflorescences (male and female) in the whole block and determine their stage of anthesis. According to this census, 15-20 inflorescences predicted to be in anthesis at the start and/or throughout each experiment were selected in each treatment such that the sticky traps were uniformly distributed over the area. The area covered by the traps was always approximatively 5.1 ha.

Sticky traps were laid above the selected (male and female) inflorescences, consisting of a Plexiglas plate, on which a parafilm was installed with glue (Figure 1,^24^). Note that only the underside of the Plexiglas plate is coated with glue, in order to capture only the weevils that leave the inflorescence.

**Figure 1:**
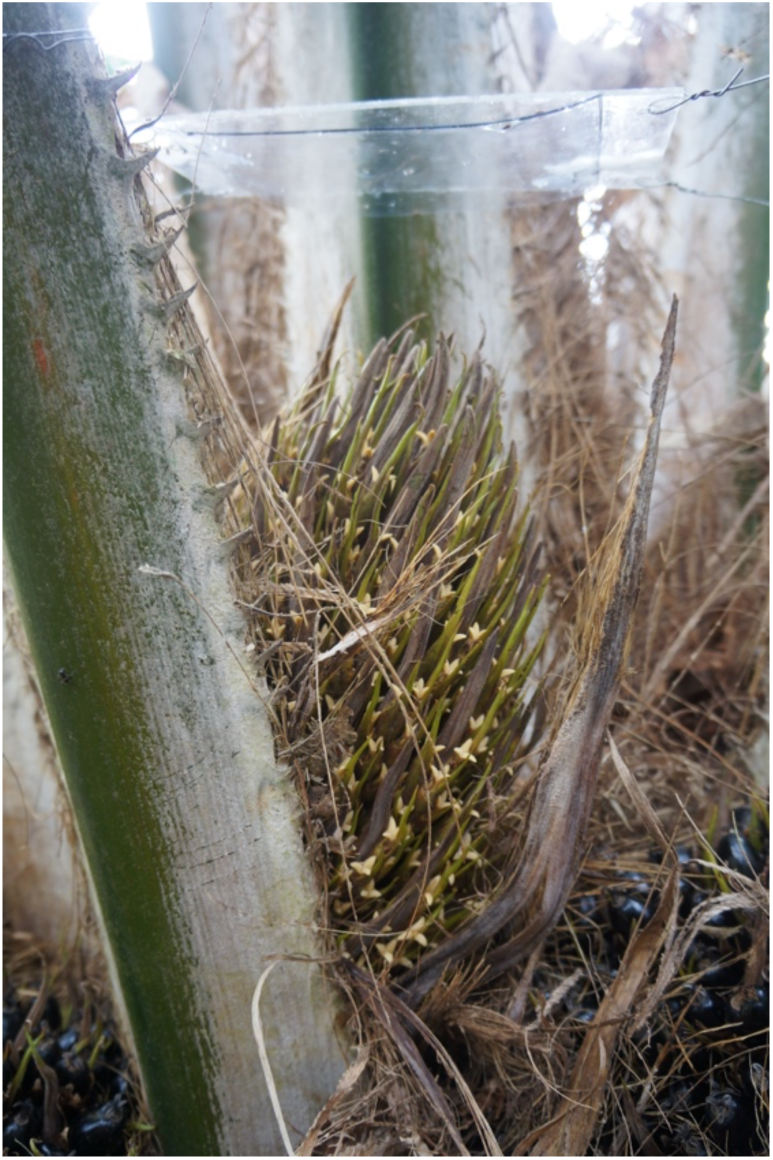
a sticky trap above a female inflorescence in anthesis.

Then, every afternoon, the traps were checked between 3:00 p.m. and 3:30 p.m.: the sticky transparent cover was removed from each trap and replaced with a new one on the same inflorescence if it has not reached the terminal anthesis stage. Then, after collection of all sticky traps in the field, each sticky trap was studied under a UV lamp to count all marked and unmarked weevils.

To give the readers an idea of the data collected during the MRR experiments, we present the raw data, by period and by year, in Figure 2.

**Figure 2:**
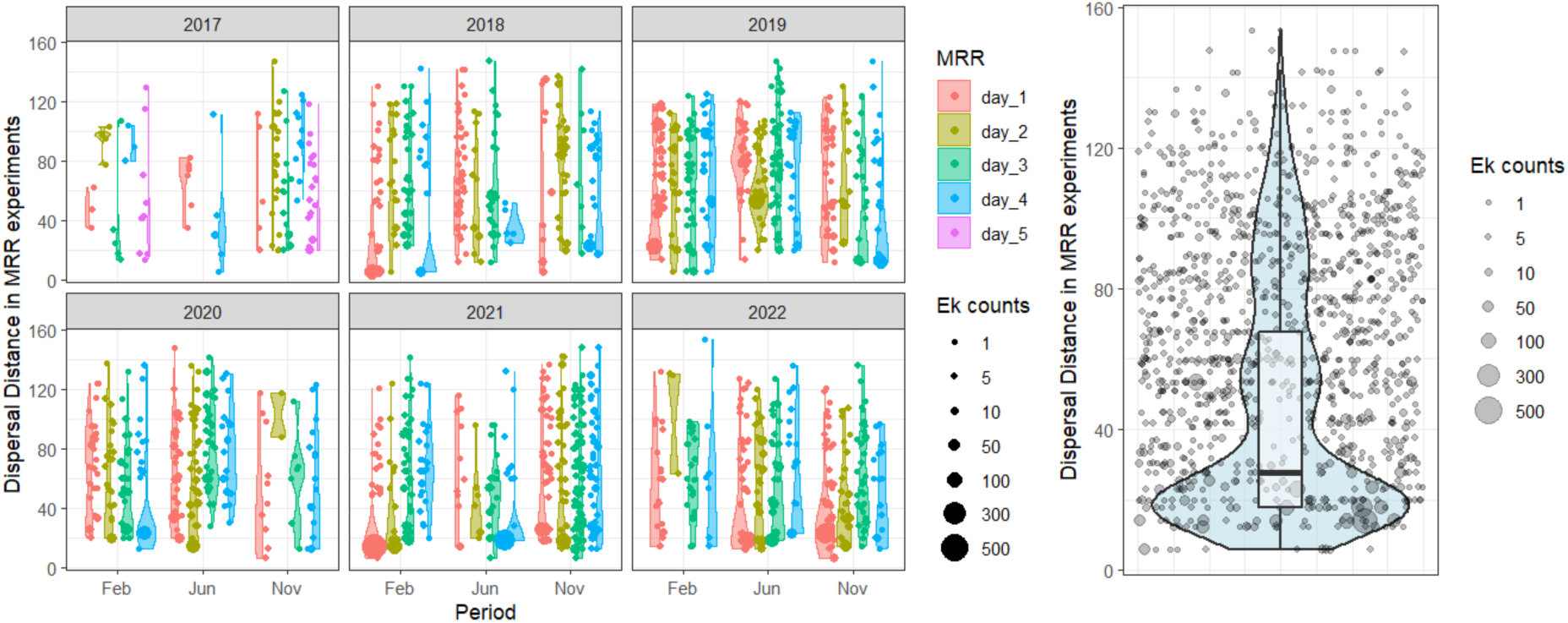
Representation of the MRR raw data. The raw dispersal distance is given in meters. Each point corresponds to a trap, with its size proportional to the number of trapped colored weevils (Ek). Density plots, figured as violin plots, and distribution figured as boxplot (only for the right panel), were overlaid. Left panel: data grouped by period, year and day of release (from day_1 to day_5). Right panel: all data together. Feb =February-March, Jun=June-July, Nov=November-December.

**Figure 3.**
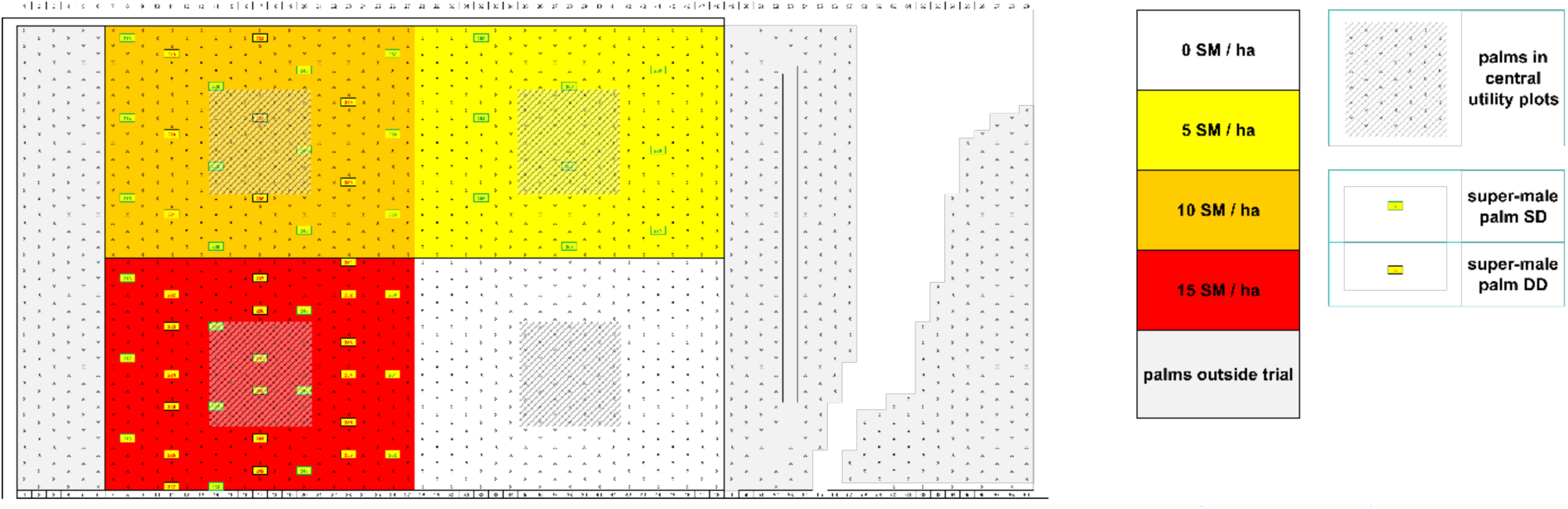
Map of the block used in this study. The map indicates the position of EXP palms (black crosses) and SM palms (yellow boxes) in each of the four plots presenting four different SM palms densities. Marked insects were released in the center of the block. This block corresponds to the block #1 located in the northern part of the super-male trial.

**Figure 4:**
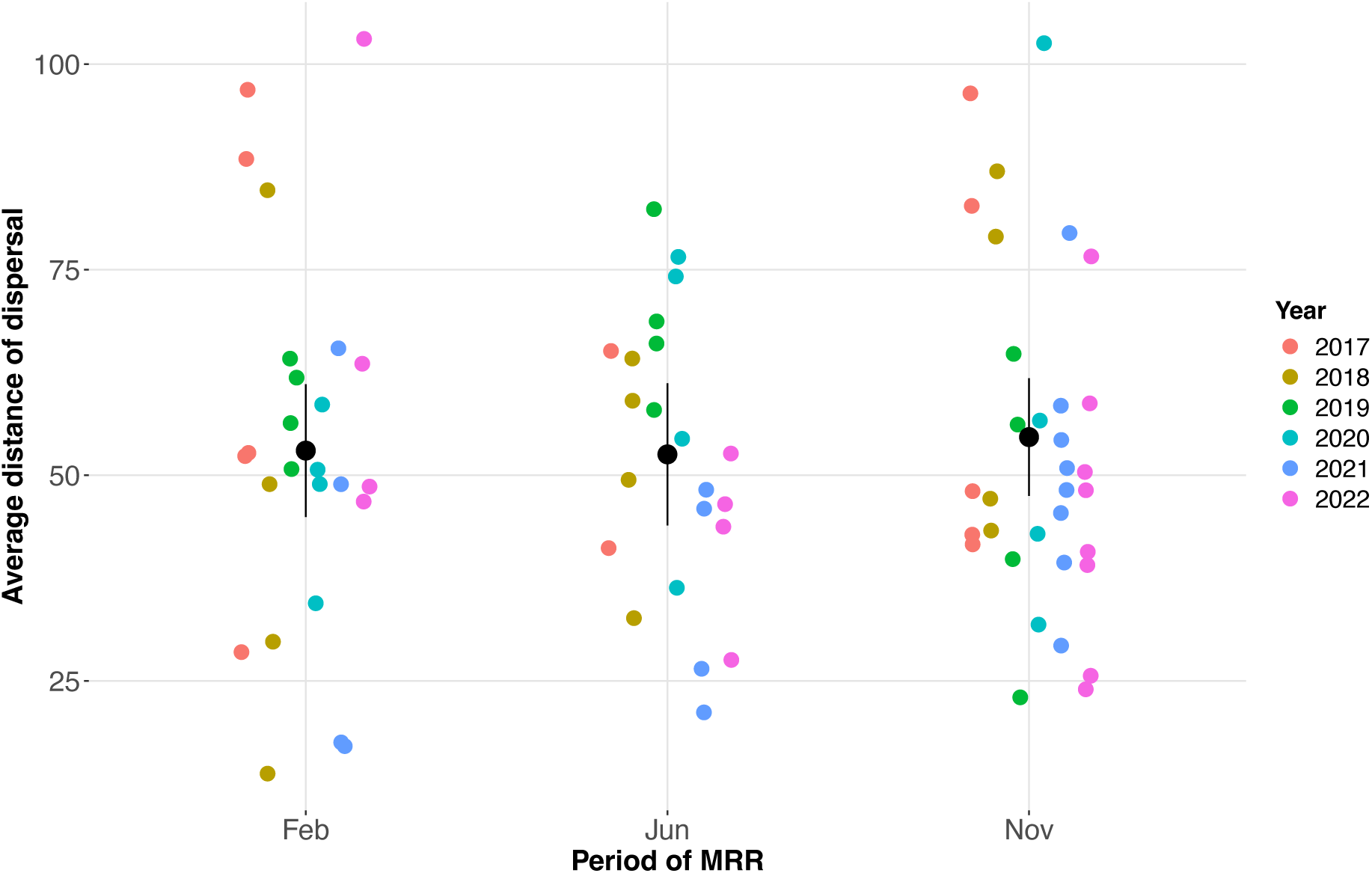
daily dispersal distance for each Mark-Release-Recapture experiment. One point corresponds to the weighted average distance in meters for one MRR experiment. Points are groups by period of the year: Feb =February-March, Jun=June-July, Nov=November-December, and colored by year.

**Figure 5:**
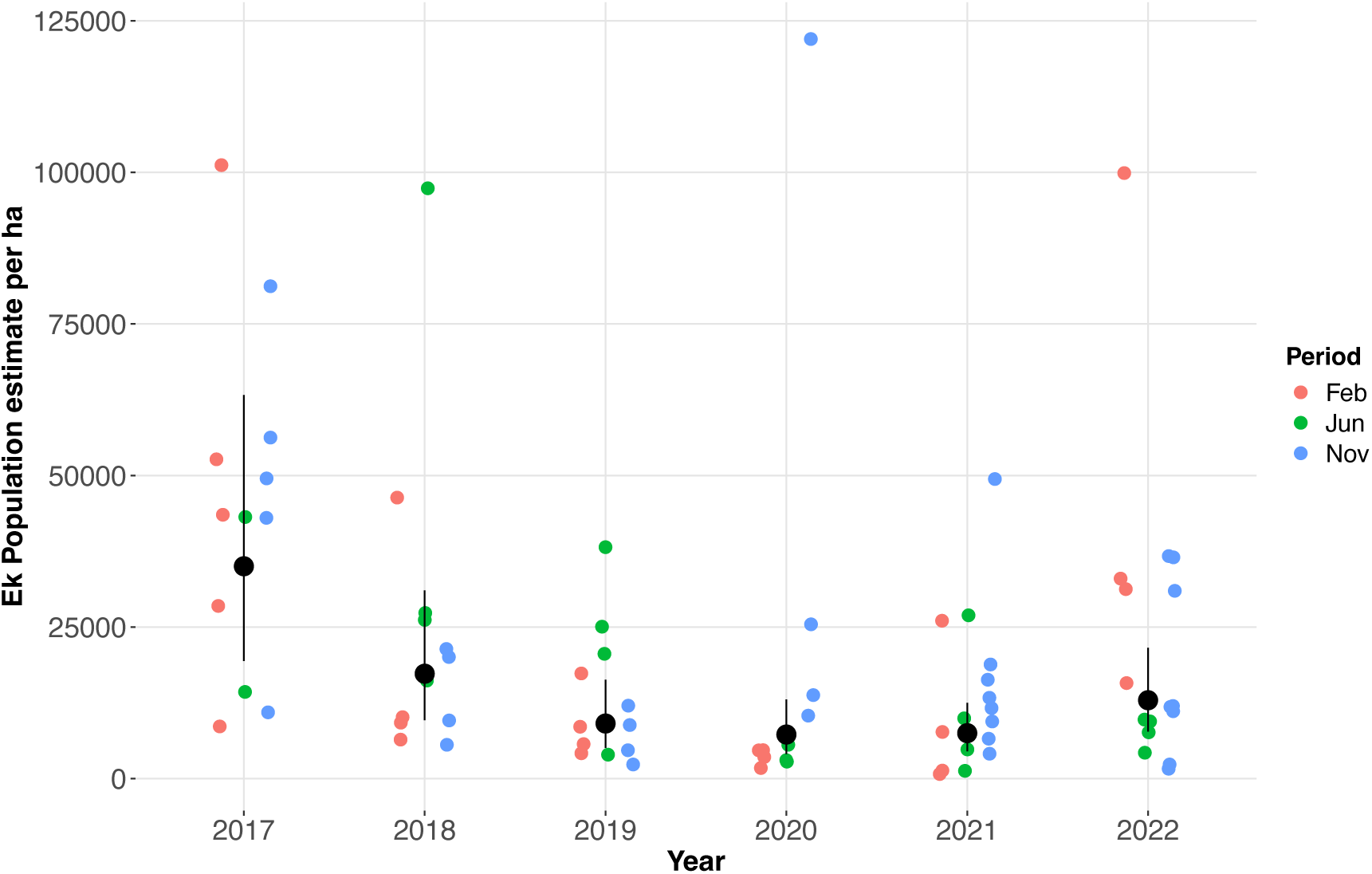
Estimates of the wild population of E. kamerunicus per hectare. Each point represents the estimate for one MRR experiment (N=82), colored by period of the year, and the means and standard deviations per year, figured in black, were obtained as estimated marginal means from a linear model with year and period as fixed effects.

For each MRR experiment (N=82), the average weighted distance of daily dispersal is estimated using the following standard formula:

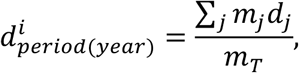

where:

- *m_T_* is the total amount of captured marked weevils during the experiment i.
- *m*_j_ is the amount of captured marked weevil in Palm j during the experiment i.
- *d*_j_ is the distance between the release point and Palm j.
- *period*(*year*) is the period of the year where the experiment i has been done.

As explained above, MRR can be used to estimate the absolute population size. Following^25^, since our daily experiments are very short (6-7 hours), we assumed that the following conditions are met: (1) the marked animals are not affected, (2) the treated animals mix well with the wild ones, (3) the capture rate of marked and wild individuals is the same, (4) sampling must be at discrete time intervals, (5) the studied population is a closed population, and (6) no births and no death occurred during the period of samplings. Then, since most of our recapture rate was low (between 0.5 and 2.5% on average, Suppl. Figure 1), for each release (N=82), the wild weevil population has been estimated according to the modified Lincoln index^26^ defined as follows:

where:

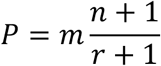

- m is the number of marked weevils released,
- n is the total number of wild weevils captured,
- r the number of marked weevils recaptured,
- P is the estimated population.

We provide a maximum of 5 population estimates per period and per year.

### Bunch analysis

During each series of experiments, 20 female inflorescences (5 per treatment) were selected for later bunch analysis (following harvest, approximately 4-5 months later), according to the level of captures (from low to high), in order to estimate the fruit set and the fruit-to-bunch ratio: see ^27,28^. To this end, each bunch was harvested and analyzed individually: after harvest, the bunch was weighted, then the different parts of bunch were separated and weighted (stalk, empty spikelets, set fruits, red parthenocarpic fruits and white parthenocarpic fruits), finally, the number of each type of fruits (set fruits, red parthenocarpic fruits and white parthenocarpic fruits) was counted.

For each analysed bunch (N=405), the fruit set was estimated with a standard formula as follows:

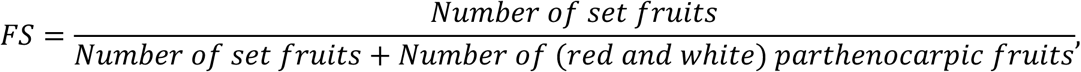

and the Fruit-To-Bunch weight ratio was estimated as follows:

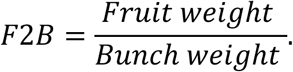

Where the fruit weight is calculated as the weight of oily fruits (set fruits and red parthenocarpic fruits).

The fruit set is considered as the best indicator to determine the pollination quality. However, the fruit set is long and tedious to estimate, while the fruit-to-bunch is faster and easier to determine. That it is why the latter is commonly used in oil palm industry as a proxy for pollination quality. Here we will consider both the fruit set and the fruit-to-bunch.

### The experimental field design: the Super-Male trial

The area of study was about 5.5 ha located within an 8-hectare block which belongs to a field trial planted in 2014 in Bangun Bandar oil palm estate (PT Socfin Indonesia, North Sumatra, Indonesia).

This trial, known as “the super-male trial”, was designed by Albert Flory (CIRAD) to study the impact of “super-male” palms on the fruit set and yield of an extremely feminine experimental oil palm cultivar. This oil palm cultivar (EXP), characterized by a very low rate of male inflorescences and a high yield, was obtained by crossing Deli palms derived from the highly feminine DA3DxDA10D population, with standard pollen used in seed production (heterotic group B) (see Figure 8). The objective was to simulate a plantation with a marked deficit in male inflorescences, a situation that can sometimes be observed when elite cultivars are planted in very favorable environmental conditions. The super-male (SM) palms used, characterized by higher ratios of male inflorescences were developed by PalmElit SAS in the frame of a dedicated breeding program. Two palms were cloned after two generations of selection for high masculinity, resulting in two SM clones named SM-1 and SM-2 (see Figure 8).

**Figure 6:**
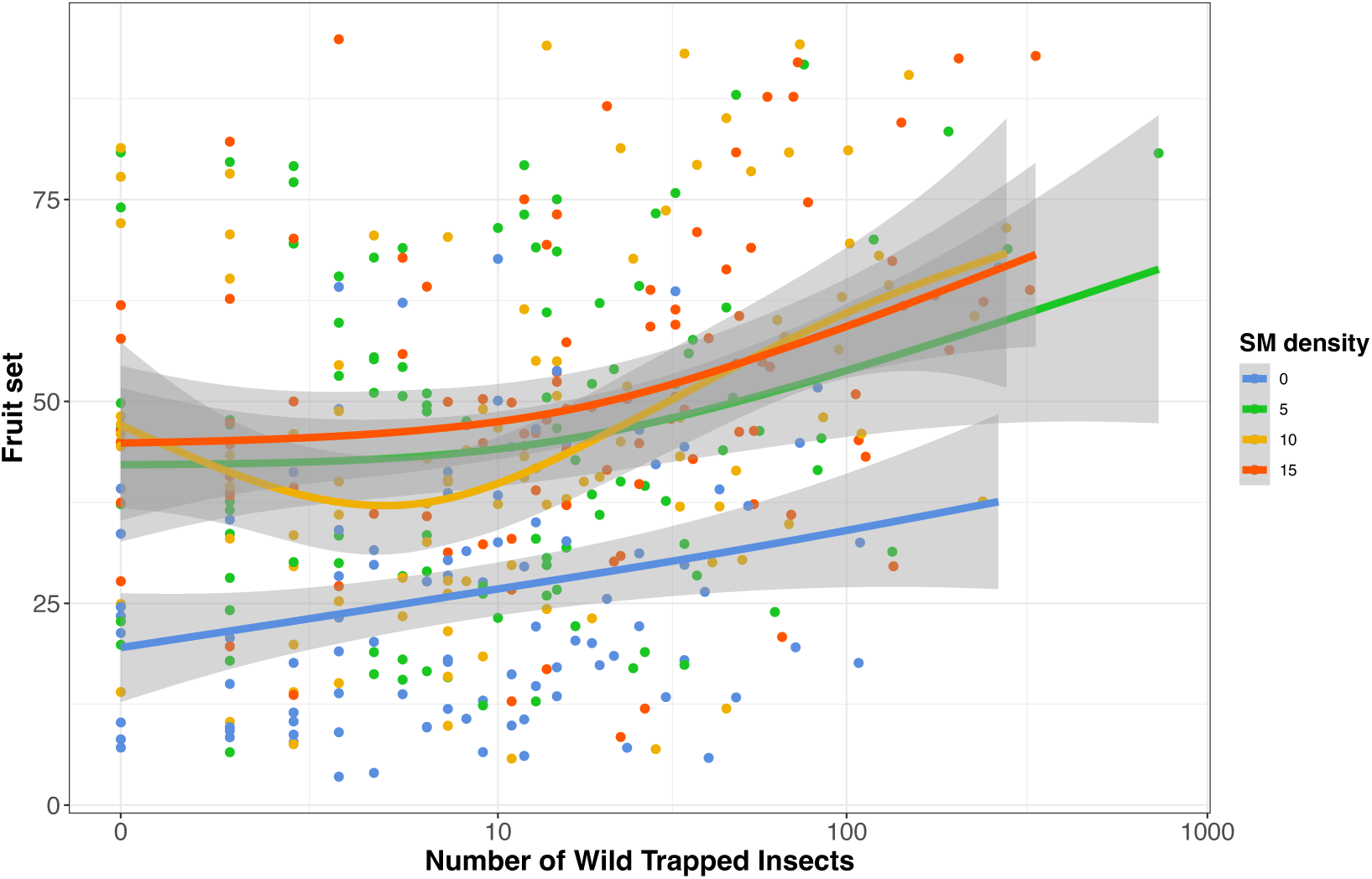
Fruit set versus number of wild trapped weevils. Each point represents an inflorescence (later a bunch, N=405). The lines correspond to smoothed conditional means with confidence intervals per SM density, as implemented in geom_smooth() function with method “gam”. Points and lines were colored according to the treatment (SM palm density).

**Figure 7:**
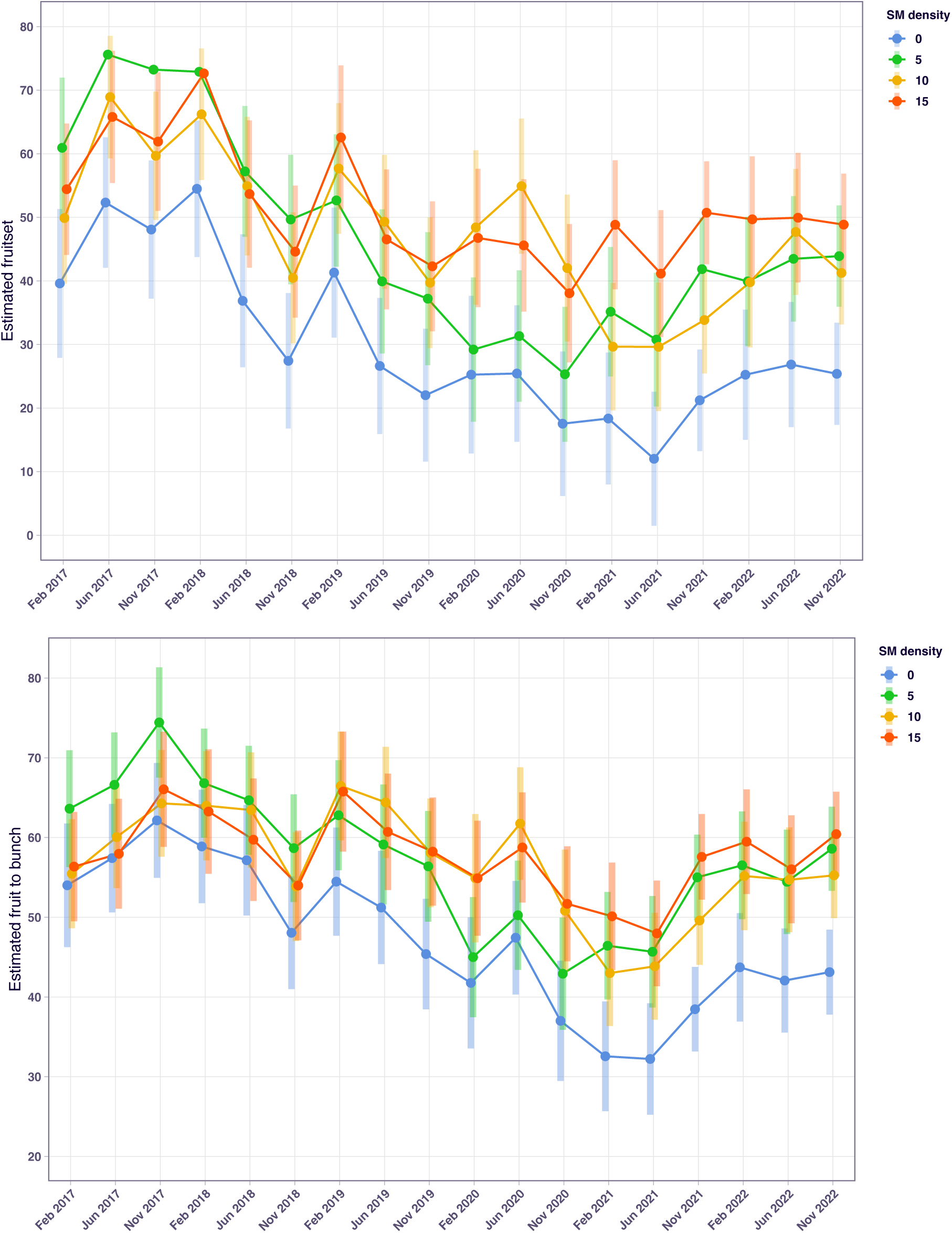
Evolution of the average fruit set (upper panel) and fruit-to-bunch (lower panel) over time, for an average recapture of 30 weevils. The period indicated in the x-axis corresponds to the period of anthesis. Error bars indicate the 95% Confidence Interval for the estimated marginal means. Points, lines and error bars were colored by SM density.

**Figure 8:**
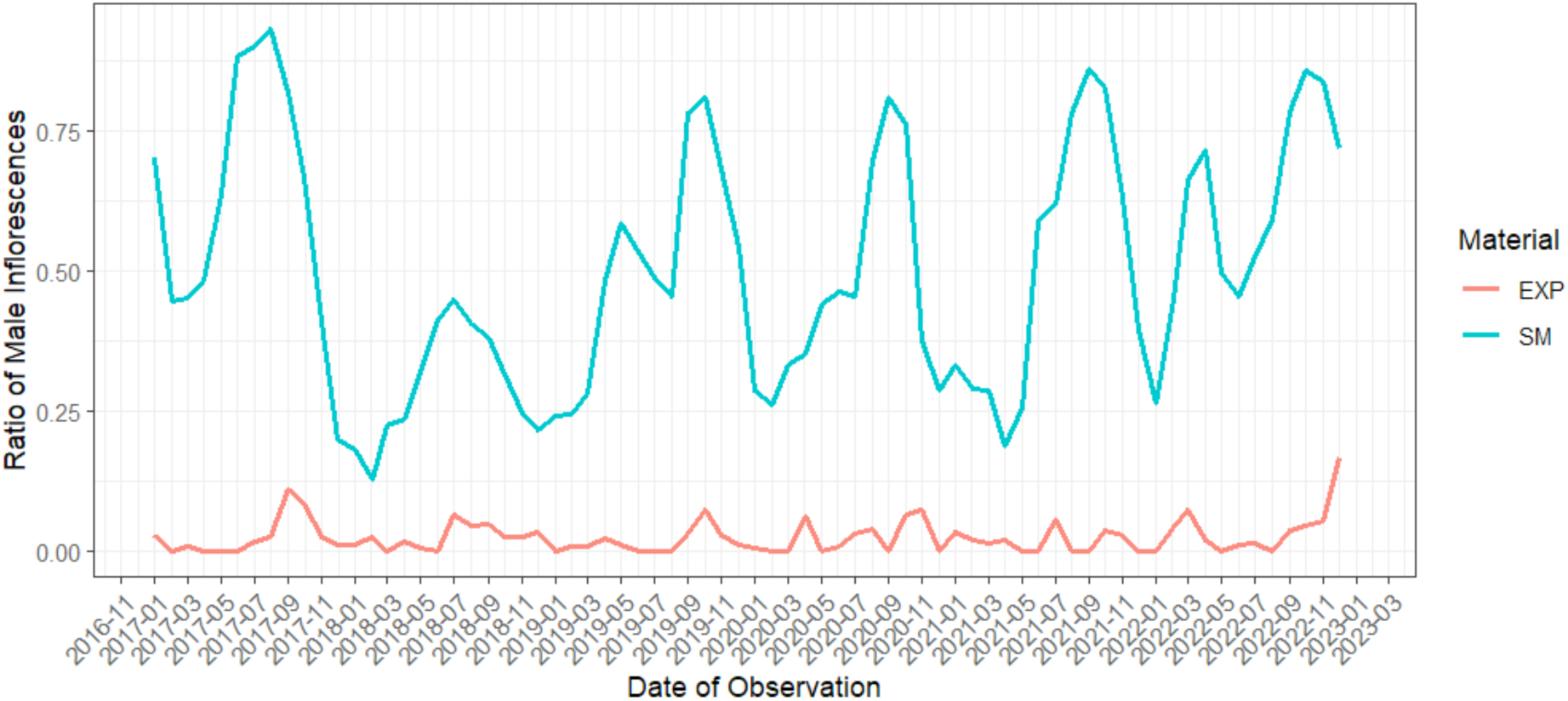
Evolution of the male inflorescences’ ratio between the Super-male (SM) material and the EXP material from February 2017 till December 2022 in block 1.

Four different treatments, with varying SM palms densities and planting strategies were tested, as explained hereafter. Male inflorescence density can be increased by increasing the density of super-male palms within productive palms, however this might at some point negatively impact the final oil yield since super-male palms will replace or compete with productive EXP palms. Indeed, super-male palms can either be planted by replacing a productive palm in the planting row (super-male palm planted in simple density (SD)) or by adding it as a supernumerary palm planted in between two existing productive palms in the row (super-male palm planted in double density (DD)). DD palms will compete with existing productive palms after a certain age, but these DD palms can be removed if they become dispensable for pollination, thus restoring a standard plantation density with limited competition among palms. Following this reasoning, the four treatments tested were as follows:

- Treatment 0: no super-male palm (143 palms/ha in total)
- Treatment 5: 5 SM palms/ha planted in SD (143 palms/ha in total)
- Treatment 10: 10 SM palms/ha with 5 planted in SD and 5 planted in DD (148 palms/ha in total)
- Treatment 15: 15 SM palms/ha with 5 planted in SD and 10 planted in DD (153 palms/ha in total)

The trial was planted in 2014 as a randomized complete block design with 8 blocks, with a total surface of 64 hectares. The details were as follows:

- Blocks 1-4 and 7-8, with EXP palms and SM-1 clone: 4 plots of ∼2.2 ha each, comprising each a central utility plot of 6.5x7 SD palms surrounded with borders of 31.5-54 m
- Blocks 5-6, with EXP palms and SM-2 clone: 4 plots of ∼1.65 ha each, comprising each a central utility plot of 5x7 SD palms surrounded with borders of 27-48 m.

Our study was conducted in block 1 (field 92A) which correspond to the replicate #1 of the super-male trial. The map of this block is presented in Figure 3**Erreur** **! Source du renvoi introuvable.**, and is representative of the other replicates. Large borders were used with the aim of isolating the central utility plots from the effects of the neighboring plots and a special care was taken for the choice of the location: the trial was planted in an area surrounded by rubber and young oil palm plantations to limit undesired effects of neighbouring pollen.

### Sex ratio measurements on productive (EXP) and super-male (SM) palms

In order to assess the sex ratio of both EXP and SM cultivars, weekly flower census were conducted in block 1 on all 59 SM palms and on a subset of 30 EXP palms selected within the central plots in each treatment (Treatment 0: 6 SD EXP palms, Treatment 5: 6 SD EXP palms, Treatment 10: 6 SD + 2 DD EXP palms, Treatment 15: 6 SD + 4 DD EXP palms). Each week, the number of new male and female inflorescences, in anthesis or post-anthesis, were recorded for each palm, and those were painted to prevent them from being counted again on the next census. A monthly sex ratio was derived for each palm, calculated as the sum of male inflorescences divided by the sum of female and male inflorescences counted during the month.

### Statistical analysis

Statistical analyses were carried out with R software (version 4.4.2, R Core Team 2024). All data are expressed as mean ± 95% asymptotic confidence interval. Depending on the type of data, we considered different types of statistical models: Linear Models (LM) and Generalized Linear Mixed Models (GLMM) were also applied. We detail the different models in the following.

Model fitting was conducted using the function “glm” from the “stats” package for LM and GLM unless otherwise stated. Model assumptions, including homoscedasticity, normality, and independence of residuals, were graphically checked. Significance of fixed effects was tested with deviance tests via the Anova function from the package “car”. When effects were significant (p < 0.05), pairwise post-hoc tests were performed using the “emmeans” function from the “emmeans” package.

### Effect of year and period on dispersal distance

To evaluate whether the average dispersal distance varied according to the year and the period of the year, a linear model (with a log variable transformation) was fitted including the effect of year, and period.

### Effect of year and period on wild EK population size

Estimated population size *P* was analyzed using a linear model including the effects of year, and period. Population estimates were log-transformed prior to analysis. Population size estimated from the fitted model for each Period x Year combination were subsequently back-transformed to the original scale for graphical representation.

### Effect of SM density, year and period on the Fruit Set

To analyze the effects of SM density, Year and Period, on the Fruit Set (FS), we considered a Generalized Linear Mixed Models (GLMM). Assuming *FS_i_*_j*k*_, the Fruit Set (FS) during period *k* and year *j* for SM density *i* to follow a beta distribution of parameter *λ_i_*_j*k*_, we used the following GLMM model with transformation *η_ijk_* = *λ_ijk_*/100.

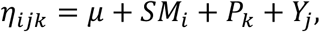

where *SM_i_*, and *P_k_* represent the fixed effects of super-male density and period, respectively, and *Y*_j_ is the random effect associated with ∀*j*, *Y*_j_∼*N*(0, *σ*^2*m*^). This model was fitted using the glmmTMB function of the R package glmmTMB ^30^ ^31^.

An Anova II test was then used to assess the significance of the fixed effects.

### Effect of SM density, year and period on the wild trapped EK

Similarly, to analyze the effects of SM density, Year and Period, on the number of unmarked trapped *E. kamerunicus* (EK), we considered a Generalized Linear Mixed Models (GLMM).

Assuming *EK_i_*_j*km*_, the number of wild trapped *E. kamerunicus* (EK) in trap *m* during period *k* and year *j* for SM density *i* to follow a Generalized Poisson distribution of parameter *λ_i_*_j*kl*_, we used the following GLMM model:

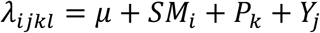

where *SM_i_*, and *P_k_* represent the fixed effects of super-male density and period, respectively, and *Y*_j_ is the random effect associated with ∀*j*, *Y*_j_∼*N*(0, *σ*^2*m*^). This model was fitted using the glmmTMB function of the R package glmmTMB.

An Anova II test was then used to assess the significance of the fixed effects.

### Effect of EK abundance, SM density, year and period on the FS and F2B

The effects of unmarked trapped *E. kamerunicus* abundance (EK), super-male density (SM), year and period on fruit set (FS) and fruit-to-bunch ratio (F2B) were analyzed separately using two similar linear models. The models included the main effects of EK, SM density, year and period, as well as all two-way interactions among these variables. Residuals were assumed to be independently and normally distributed. The significance of model terms was assessed using analysis of variance (ANOVA).

Since we expect the density of SM palms to affect the number of male inflorescences and consequently the pollen availability and transport by the weevils, we derived estimated marginal means adjusted from the effect of EK. More precisely, to compare SM density treatments independently of fluctuations in pollinator abundance, estimated marginal means for each SM x year x period combination were calculated for a fixed value of EK=30, corresponding to the overall average number of unmarked trapped weevils recorded across all MRR experiments conducted between February 2017 and December 2022 (N=82).

These estimated marginal means and their associated 95% confidence intervals were then used to visualize the temporal evolution of FS and F2B under this standardized pollinator abundance.

## Results

### Estimates of daily dispersal distance

Across 82 MRR experiments conducted over six years at three different periods in the year, we estimated daily dispersal distances for the released marked *E. kamerunicus* based on all traps positioned within the oil palm plot. As shown in , most weevils were recaptured within 40 m from the release spot, but some weevils were captured at distances above 150 m, suggesting that at least some insects can travel over long distances within one day. For each MRR experiment (N=82), we derived an estimate of the average daily dispersal distance (Figure 4).

As can be seen in Figure 4, the daily average dispersal distance can drastically vary at a given period of the year. However, the average dispersal per period is almost similar: 53.0±8.1 m in February, 52.5±8.7 m in June, and 54.6 ±7.2 m in November. The maximal (minimal) average dispersal in February and in November is 103 m (13.7 m and 23 m respectively), and 82.4 (21.2) m in June. Using a linear model, and a type II Anova analysis, we show that the period and the year have statistically no significant impact on the average dispersal distance (p>0.2381). From this result, we conclude that under the study conditions, the average daily dispersal distance isn’t clearly influenced by the year and the period of the year, with a mean value of 53,37±8.0 m.

### Estimates of the weevils’ population

For each MRR experiment (N=82), we also estimated the average population size of wild *E. kamerunicus* per ha (Figure 5). Using a linear model and a type II Anova analysis, we show that only the year has a significant impact on the average population size (p< 0.001502), while the period has no significant impact (p>0.5136).

Then, using marginal means derived from the previous linear model we derive Figure 5.

The annual average of the population, expressed as the number of individuals/ha, ranged from 6 682 (in 2021, 7 years after planting) to 32 890 (in 2017, 3 years after planting).

Unexpectedly, an overall decrease of population density was observed from 2017 till 2020, with a stabilization in 2021, and then an increase in 2022.

In the next section, we assessed whether this population fluctuation can impact the fruit set and fruit-to-bunch ratio, in relationship with the SM palm density.

### Impact of the SM planting design on the fruit set (FS), the fruit-to-bunch ratio (F2B), and the number of trapped wild *E. kamerunicus*

In the super-male field trial, the main objective was to observe the impact of the SM palm density on the fruit set (FS) and the fruit-to-bunch (F2B) ratio. Our MRR experiments provided additional information to dissect the link between the density of super-male palms, the number of wild trapped *E. kamerunicus*, and the evolution of the fruit set over time. To do so, a subsample (N=405) of the inflorescences monitored in MRR were selected for a subsequent measurement of fruit set and fruit-to-bunch. A representation of the fruit set versus the number of trapped wild EK on the selected inflorescences already suggested that there might be a positive correlation between both variables, and that this relationship depends on the SM palm density (Figure 6).

We first analyzed the effect of the SM density on the fruit set using a simple model. Using a GLMM model and a likelihood ratio test, we show that the SM density has a highly significant positive impact on the fruit set (*p* < 2*e*^−16^), while the period has only a small significant impact (*p* < 0.03254). A first pairwise analysis shows that the fruit set at density 0 is significantly smaller than all other densities (*p* < 0.0001). The fruit set in densities 5, 10, and 15, are statistically similar (*p* = 0.0529 for density 5 vs density 15, *p = 0.9937* for density 5 vs density 10*, and p = 0.1029 for density 10 vs density 15). Similarly, another pairwise analysis shows that the period has a moderate impact on the fruit* s*et: the lowest* f*ruit* s*et is reached in November and the highest* in June, February being at intermediate, but without statistically significant difference with June (p *= 0.9745*) and November (p *= 0.0912*).

We then tried to identify which pollination component is impacted by the SM density. Firstly, we analyzed the effect of the SM density on the number of trapped wild EK per inflorescence. Using a GLMM model, a likelihood ratio test showed that the period has no significant effect on the number of wild trapped insects per female inflorescence (*p*=0.5648), while the SM density has a positive impact on the number of wild trapped weevils (*p*=0.04687). However, a pairwise analysis showed that there is only a significant difference between the wild weevils trapped in density 15 and those trapped in density 0 (*p*=0.0461).

For all other pairwise comparisons, there were no significant differences. These results suggest that the weevil population is higher in plots containing SM palms, a factor which likely contributes to the observed increase in fruit set.

Secondly, we analyzed whether the SM density might impact other pollination components, independently of the number of trapped wild weevils. Using the MRR data as well as the fruit set and the fruit-to-bunch data, we used a statistical model that allows to derive the time variation of the fruit set and the fruit-to-bunch along the 6 years, for a fixed average amount of wild trapped weevils. From this model, for the fruit set, we derived the analysis of variance presented in **Erreur ! Source du renvoi introuvable.**, and estimated marginal means presented in Figure 7.

From Table 1, we deduce that some variables have a highly significant impact on fruit set: the number of trapped weevils (EK), the SM Density, the Year, and the interaction between the Period and the Year. The Period and the interaction between the SM Density and the Year also have an effect on fruit set, although less significant.

**Table 1:** Summary statistics of the linear model used to study the effect of EK, SM density, Year and Period on fruit set.

| Response: FS | Df | Sum Sq | Mean Sq | F value | Pr(>F) |
| --- | --- | --- | --- | --- | --- |
| EK | 1 | 19918.2996 | 19918.2996 | 86.3063272 | 0.0000000 |
| SM Density | 3 | 30451.2919 | 10150.4306 | 43.9819866 | 0.0000000 |
| Period | 2 | 2612.2981 | 1306.1490 | 5.6595658 | 0.0038115 |
| Year | 5 | 32175.2729 | 6435.0546 | 27.8831997 | 0.0000000 |
| EK:SM Density | 3 | 1628.9404 | 542.9801 | 2.3527420 | 0.0719685 |
| EK:Period | 2 | 731.9598 | 365.9799 | 1.5857971 | 0.2062565 |
| EK:Year | 5 | 1643.2619 | 328.6524 | 1.4240563 | 0.2148445 |
| SM Density:Period | 6 | 890.0229 | 148.3372 | 0.6427474 | 0.6959872 |
| SM Density:Year | 15 | 6806.6095 | 453.7740 | 1.9662102 | 0.0168915 |
| Period:Year | 10 | 11899.0231 | 1189.9023 | 5.1558667 | 0.0000005 |
| Residuals | 349 | 80544.3447 | 230.7861 | - | - |

The estimated means, for a fixed number of visiting weevils, shows that treatment 0 provides the lowest fruit set, compared with treatments with SM palms, with treatment 15 showing most often the highest fruit set. Overall, these results show that the SM density has a positive impact on the fruit set, independently of the number of trapped wild weevils. This likely illustrates that there is more viable pollen available in plots containing SM palms.

Then, using the same type of model, we derived estimates for the fruit-to-bunch over time (Figure 7, lower panel), which follow the same tendency as the fruit set. Thus, the density of SM also positively impacts the fruit-to-bunch, which is a key yield component. Interestingly, the estimates for both fruit set and fruit-to-bunch showed similar variations over time, independently of the SM density, with a decreasing trend over the years. Since these estimates were obtained for a fixed number of visiting weevils, these variations could be due to environmental factors other than fluctuations in *E. kamerunicus* population.

Finally, to better understand how SM density affects pollen availability, we studied the male inflorescence production of SM palms, in comparison with that of EXP palms. In block 1, phenology of male and female inflorescences was conducted weekly for all SM palms (59 palms) and a subset (30) of the EXP palms such that their inflorescence’s dynamics is known all along the MRR period. Using these data, we calculated the ratio of male inflorescences versus the total number of inflorescences per month and per cultivar (Figure 8).

This result shows that the ratio of male inflorescences of the SM palms is always higher than for EXP palms. However, this ratio shows some seasonal variations with an important peak from June to October every year for the SM palms, except in 2018. The variations of sex ratio over time likely contributes to the variations of fruit set and the fruit-to-bunch observed in Figure 7. Over the whole period, SM palms have produced, on average, 61.0% male inflorescences, while EXP palms, only 3.04%.

Collectively, these results show that SM density positively influences fruit set and fruit to bunch, and that this effect is linked with an increase of both the number of *E. kamerunicus* visiting the female inflorescences, and of the amount of viable pollen available in the plot. Consistently, SM palms demonstrates a good potential as pollinator palm, characterized by a very high rate of male inflorescences (61.0% versus 3.04% for EXP palms), despite some seasonal variations.

## Discussion

### *E. kamerunicus* dispersal dynamics

Generally, Mark-Release-Recapture experiments are complex and difficult to conduct in the field. However, in addition to monitoring traps that might (only) give relative information about the time evolution of pollinators abundance, MRR provides various useful information, like the average dispersal distance and also estimates of the absolute population that are useful factors to monitor in particular when fruit set variations occur. Average dispersal distance is also a critical factor for optimizing planting designs when these include pollinator palms such as the SM palms presented here.

Our study showed that the average dispersal distance is 52.843 m., with a maximum of 143.78 m., which is higher than what could have been expected. This highlights the importance of using large borders (>50 m) for field trials involving different pollination treatments. Interestingly, the average dipersal distance didn’t show any significant variation among years and months, and thus seems to be overall stable in the study conditions.

We also show population estimates of 6700 to 32 800 individuals, with some unexplained variations along the years. It is well known that large population is a requirement to reach a good level of pollination, i.e. with a fruit set above 70%^1^. However, other factors are critical, especially pollen quality and quantity, and these can be influenced by the planted material/cultivar as well as environmental factors such as the weather.

### Impact of the super-male palms

The originality of this work was to conduct the MRR experiments within a unique field design, the “super-male trial”, to study the impact of several super-male palm densities (0/5/10/15 SM/ha) on key parameters related to the fruit set. As expected, the SM palms show a higher ratio of male inflorescences (61%), as compared to the EXP palms (3%), meaning that the overall density of male inflorescences in the field increases with the density of SM palms. Accordingly, we show that there is a significant increase in fruit set and fruit-to-bunch ratio in plots containing SM palms, versus the plot without SM palm. Thanks to the MRR experiments, we could show that this positive effect is likely due to a joint increase in the number of *E. kamerunicus* visits and in pollen availability, keeping in mind that these two parameters are likely correlated with each other. We can hypothesize that the number of visiting *E. kamerunicus* increases because a higher density of male inflorescence will attract more weevils and at the same time allow the development of a larger weevil population.

Although we don’t show here the impact on final oil yield, we can assume that the increase in fruit set and fruit-to-bunch translates into a yield increase, since the fruit set in the plot without SM palm remains largely below the optimum of ∼70%. Our results thus show very promising results regarding the use of SM palms to increase oil yield in plantations where there is a deficit in male inflorescences, especially at the young age. However, there is a need to conduct additional studies to confirm these results at a larger scale and to assess the economical return of the different planting scenarios over longer periods. Of note, fruit set remained relatively low over the whole period of study (e.g. 70% of bunches in the treatment 15 had fruit set values below 60%). This raises several questions such as the impact of other environmental factors that could negatively affect fruit set and whether the male inflorescence density in the trial remained a limiting factor for pollen production and/or maintenance of the *E. kamerunicus* population.

### Towards optimized planting designs

In ^32^, we developed and compared a macroscopic mathematical model and a microscopic individual based model to derive estimate of the weevil’s population needed to reach a fruit set of 70%. The new results obtained in this study will help to develop a more complex mathematical model with two planting materials to determine the best/optimal super-male planting density to insure the best fruit set along the years. These in-silico results may help planters in their choice of SM plantation density and spatial layout inside each plot. Such modeling approach will be essential to determine the optimal planting parameters since field testing is very costly and lengthy.

Still, additional results would improve the outputs. For instance, the temporal evolution of the ratio between viable pollen and total available pollen. This is a rather important issue because it may also impact on the fruit set, even if the weevils’ population is sufficiently large to guarantee a good pollen dispersal.

Since increasing the pollinator palm density results in competition with productive palms planted in the same plot, fruit set alone isn’t sufficient to account for the final economic return per hectare. Thus, a more comprehensive study that takes into account the yield together with the operating costs will be necessary to determine the optimal planting design. The final objective is to identify the sweet spot that balances pollinator palms density and productive palms density. Planting strategies where all or some pollinator palms are uprooted after a certain age should also be studied.

## Conclusion

Using extensive mark-release-recapture experiments conducted over several years in a unique pollination field trial planted in North Sumatra, we report estimates of key parameters for the pollinating weevil, *E. kamerunicus*, such as population size, average and maximum daily dispersal distances. These parameters will be useful for modelling the interactions between oil palm and its pollinator weevil, to support the optimization of planting designs and pollination processes.

Furthermore, our MRR experiments, combined with the analysis of a large number of monitored bunches allow us to study more finely the impact of the super-male palms planting designs on fruit set and fruit-to-bunch ratio. Our results show a positive effect of super-male palms on fruit set and fruit-to-bunch ratio, consistently with their high rate of male inflorescences. We provide evidence that this positive effect results from a double positive effect on *E. kamerunicus* population size and on pollen availability. Still, a finer analysis will be required to determine the optimal plantation design for super-male palms, in terms of density per hectare and proportion to plant in single and/or double density, in order to achieve a balance between competition effects and fruit set improvement.

## Data

All data will be available at: https://doi.org/10.18167/DVN1/ORYKYQ

## Acknowledgements

All authors acknowledge the technical and financial support of PALMELIT SAS and of PT SOCFIN INDONESIA where all experiments were conducted.

## Author Contribution

YD and LBO conceived and designed research. AL produced the SM palms. YD, LBO, HR, CM, IS, and DA conducted experiments. YD, AD, and FJ performed the statistical analyses. YD and FJ wrote the original draft. All authors read and approved the manuscript.

## Supplementary material

**Supplementary Figure 1:**
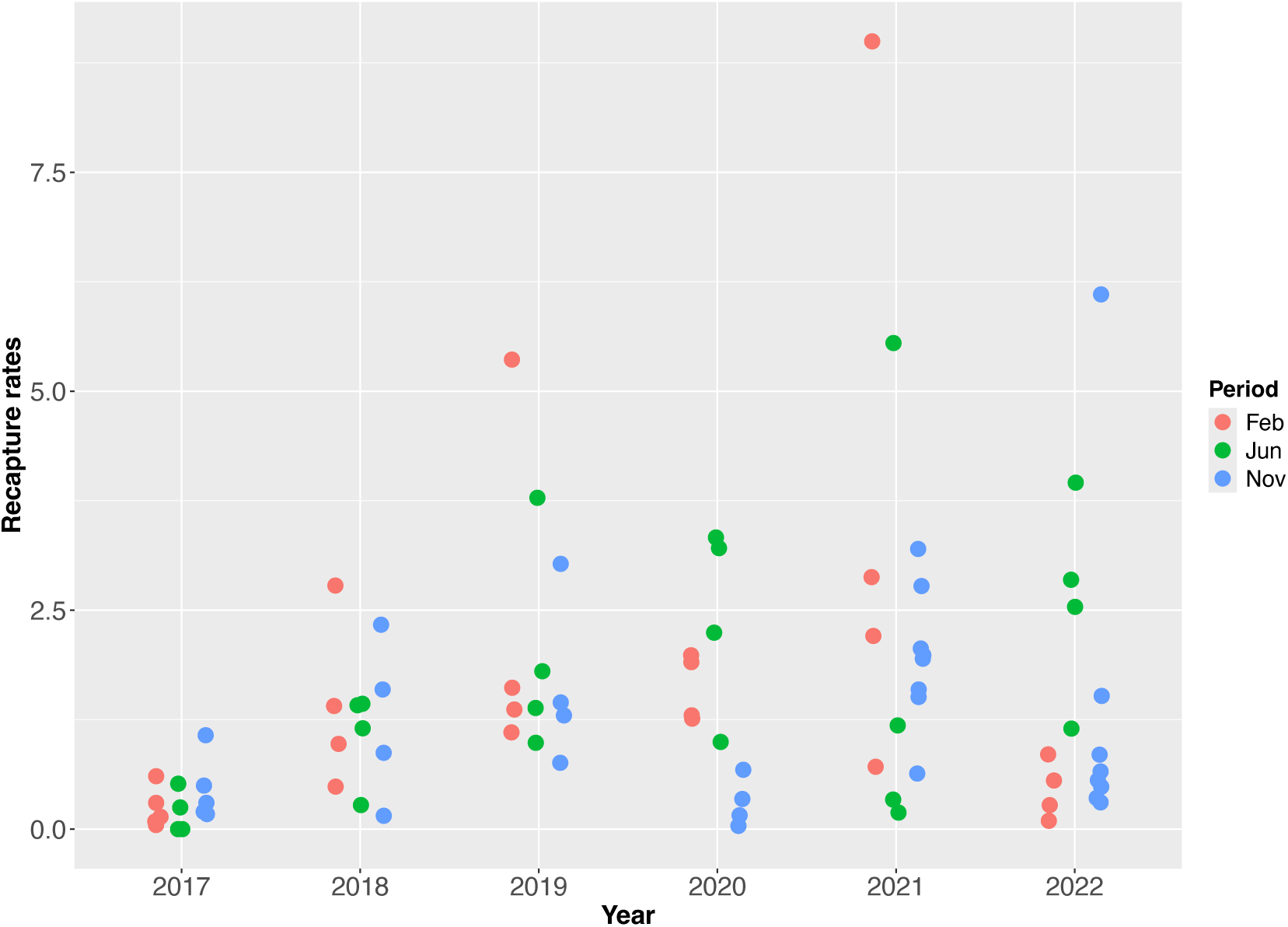
Recapture rates (in %) for each MRR experiment from 2017 till 2022 for each Period (February in red, June in green, and November in blue (N=82).

